# Characterization and automated classification of sentences in the biomedical literature: a case study for biocuration of gene expression and protein kinase activity

**DOI:** 10.1101/2025.01.06.631539

**Authors:** Daniela Raciti, Kimberly M. Van Auken, Valerio Arnaboldi, Christopher J. Tabone, Hans-Michael Muller, Paul W. Sternberg

## Abstract

Biological knowledgebases are essential resources for biomedical researchers, providing ready access to gene function and genomic data. Professional, manual curation of knowledgebases, however, is labor-intensive and thus high-performing machine learning methods that improve biocuration efficiency are needed. Here we report on sentence-level classification to identify biocuration-relevant sentences in the full text of published references for two gene function data types: gene expression and protein kinase activity. We performed a detailed characterization of sentences from references in the WormBase bibliography and used this characterization to define three tasks for classifying sentences as either 1) fully curatable, 2) fully and partially curatable, or 3) all language-related. We evaluated various machine learning (ML) models applied to these tasks and found that GPT and BioBERT achieve the highest average performance, resulting in F1 performance scores ranging from 0.89 to 0.99 depending upon the task. Our findings demonstrate the feasibility of extracting biocuration-relevant sentences from full text. Integrating these models into professional biocuration workflows, such as those used by the Alliance of Genome Resources and the ACKnowledge community curation platform, might well facilitate efficient and accurate annotation of the biomedical literature.

## Introduction

Biological knowledgebases, hereafter referred to as knowledgebases, are a key resource for biomedical researchers, affording them ready access to gene function and genomic data (1). Largely populated by professional biocurators, knowledgebases adhere to strict data models and standards in accordance with FAIR principles (Findable, Accessible, Interoperable, and Reusable) (2). An important aspect of FAIR biocuration is preserving the provenance of biological knowledge, which typically involves a manual process of identifying relevant published data, translating that data into models using standardized identifiers and ontologies, and associating the curated data with the original peer-reviewed reference (3). While manual biocuration provides knowledgebases with detailed, high-quality data, the process is labor-intensive, and the increasing number of references, as well as the complexity of data within those references, has necessitated development of semi-automated approaches to ensure that experimental findings are incorporated into knowledgebases in a timely manner. Additionally, recent funding cuts to manual biocuration projects emphasize the urgent need to focus on developing biocuration methods that are more efficient, yet still of uncompromising quality (4–6).

Two main approaches to this curation challenge adopted by biological knowledgebases are: 1) artificial intelligence (AI) and machine learning (ML) methods, and 2) community curation. AI and ML methods have been successfully employed at several stages of the curation process including identifying: 1) curation-relevant references, 2) specific data types within those references, 3) entities such as genes, alleles, and strains used in experiments, and 4) sentences for fact extraction (for illustrative examples see 7-17). In addition, some knowledgebases also rely on authors for curatorial assistance, as these subject matter experts can, with guidance from professional biocurators, provide high quality curation of their published references (18–23).

The ACKnowledge (Author Curation to Knowledgebase) project combines the strengths of state-of-the-art ML approaches with author expertise in a community curation pipeline (20). Initially implemented at WormBase (24), the pipeline automatically extracts biological entities such as genes and strains from the full text of recently published scientific articles and predicts the presence of nine curation-relevant data types such as gene expression and catalytic activity. Authors are then presented with a web form containing the extracted information where they can validate and submit data to WormBase.

While the current ACKnowledge system allows authors to contribute valuable information on entities and data types, our ultimate goal is to actively engage them in submitting more comprehensive curation, i.e. fact extraction statements such as “gene A is expressed in cell B” or “protein C phosphorylates protein D”. To this end, we are furthering our approach by developing methods that combine ML with community curation to identify relevant text for fact extraction. The identified text will then be presented to authors to assist them in curation. While we are developing these methods to assist community curation, they will also be invaluable in professional biocuration workflows.

Relevant work in the literature on text classification include the work of Shatkay et al. (25), who introduced a multi-dimensional framework for annotating biomedical sentences, incorporating scientific focus, generic nature, and statement polarity, which paved the way for further research in the field. Other methods have focused on classifying sentences in PubMed abstracts into broad semantic and rhetorical categories, such as Introduction/Background, Methods, Results, and Conclusions, which further aid in text mining tasks. These methods vary widely, including Naive Bayes models, support vector machines, Hidden Markov models, Conditional Random Fields (CRFs), and other methods based on more sophisticated feature engineering (for examples, see 26,27).

Additional work in sentence identification was performed as part of the BioCreative IV GO task in which curators collected two types of evidence sentences, summary and experimental, to support GO annotations (28). Using these evidence sentences as training data, participating teams attempted to locate GO-relevant sentences from full-text articles given relevant gene information in the text retrieval task (29). The results of this task, however, indicated that much work still needed to be done to identify curation-relevant sentences, as the F1 scores for identifying exact or overlapping evidence sentences were only 0.270 and 0.387, respectively.

The BioCreative V BioC track (30) aimed to develop a collaborative biocurator assistant system for BioGRID using the BioC XML format to manage text, annotations, and relations. The system analyzed full-text articles, prioritizing sentences reporting experimental methods and mentions of protein-protein and genetic interactions. BioGRID curators evaluated the system, giving positive feedback on its usability and design, though they suggested improving passage prediction accuracy, with F1 scores ranging from 0.65 to 0.8, depending on the task.

Recent advances in sentence and text classification have further enhanced the capabilities of biocuration. BERT (Bidirectional Encoder Representations from Transformers) is a state-of-the-art language representation model developed by Devlin et al. (31). This model leverages a transformer architecture to pre-train on a large corpus of text, enabling it to capture deep bidirectional context. BERT proved to be a robust model for text classification, outperforming previous solutions. Models such as BioWordVec (32) and BioSentVec (33), are able to generate word and sentence embeddings specifically for biomedical texts and demonstrated considerable promise. Each of these models was trained on over 30 million documents from PubMed and clinical notes from the MIMIC-III Clinical Database, providing robust embeddings for word and sentence pair similarity tasks. These embeddings can be easily combined with more traditional ML models such as SVMs and Logistic regression to perform sentence classification with lightweight models compared to those based on BERT and GPT. BioBERT (Bidirectional Encoder Representations from Transformers for Biomedical Text Mining), a domain-specific language representation model pre-trained on large-scale biomedical corpora, has outperformed BERT and previous state-of-the-art models in a range of biomedical text mining tasks, including named entity recognition, relation extraction, question answering, and text classification (34).

Most recently, Generative Pre-trained Transformer (GPT) models (35) have demonstrated remarkable capabilities primarily in text generation but also in classification tasks. These models leverage extensive pretraining on diverse datasets, making them versatile for various NLP tasks, including sentence classification (36).

In this study, we focus on sentence-level classification for identifying curatable information from full text articles. This approach simplifies the identification process by treating sentences as discrete units of text that convey a statement and are complete in themselves, rather than attempting to extract relevant information from the entire body of scientific articles. By focusing on individual sentences, we hope to more effectively isolate and analyze specific pieces of information, which may enhance the accuracy and efficiency of our curation efforts. This method also reduces the complexity associated with parsing and interpreting longer, more intricate texts, making it a more manageable and scalable solution for biocuration tasks. In addition, identifying curation-relevant sentences can facilitate manual validation of fact and entity extraction by highlighting the most critical information needed for curation.

To achieve our aims, we first performed a detailed characterization of the types of sentences relevant for curation of gene expression and protein kinase activity, two key data types curated at WormBase. We then grouped these sentences into classes for the purpose of performing biocuration tasks of increasing specificity. Using the identified classes, we trained ML and AI models to automatically classify relevant sentences. GPT and BioBERT models demonstrate superior average performance, with F1 scores ranging from 0.89 to 0.99 depending upon the task.

Our findings demonstrate the feasibility of extracting biocuration-relevant sentences from published scientific references. Integrating these sentence classifiers into the ACKnowledge, as well as professional biocuration platforms, will enhance community and professional biocuration efforts, by focusing work on high-value curation statements, thus maximizing curation efforts.

## Methods

### Full text data collection and preprocessing

#### Source of data

The full text of references was obtained from the Alliance of Genome Resources (The Alliance), which now serves as the primary literature data source for WormBase curation and ACKnowledge processing. As part of shared infrastructure, the Alliance has developed the Alliance Bibliographic Central (ABC), a platform to manage references including their full text files, which are primarily stored as PDFs (37). We accessed these PDFs using the ABC API and converted them to plain text with GROBID (https://github.com/kermitt2/grobid), a state of the art machine learning-based library for parsing structured file formats and converting them to standard XML, that is also able to identify and extract sentences. We applied XML manipulation to obtain the list of plain text sentences from each article sourcing them from all sections except the cited references. The high variability in templates across different journals and articles makes it challenging to consistently identify specific sections beyond the cited references, which are always correctly removed by GROBID.

#### Cluster analysis

We used visual techniques to assess the similarity between sentences and assess if sentences from the same classes (following the definitions provided below) were clustered together. Specifically, we embedded text sentences using BioSentVec and then we applied Uniform Manifold Approximation and Projection (UMAP, 38), a dimensionality reduction technique that allowed us to visualize sentence representations in two dimensions.

### Sentence datasets acquisition

#### Overview

To create a dataset for sentence classification, we collected both data type-relevant sentences and non-relevant, or “negative”, sentences. Negative sentences do not contain language pertinent to the data types of interest. We identified sentences via both random sampling and through manual selection from references in the WormBase bibliography. We included manually selected sentences because random sampling alone did not yield a sufficient number of data type-relevant sentences for our analysis.

Data type-relevant sentences either described experimental results for a given data type, described related information such as experimental details, or contained language related to the data type but not directly relevant to the curation of that reference. A more detailed description of the differences between data type-relevant sentences is provided below.

#### Random sampling

We collected a set of 1000 randomly selected sentences sourced from 15 manually validated positive and 15 manually validated negative references for the data types. These sentences were then manually reviewed and labeled by a curator according to the criteria outlined below.

#### Manually extracted data type-relevant sentences

We collected 589 and 494 data type-relevant sentences from 115 and 51 references, respectively, for gene expression and protein kinase activity (Supplementary Table 1). For each of the data type-relevant sentences, curators characterized the type and amount of information it contained (see the Results section for more examples) and collected relevant metadata, such as the section of the reference where the sentence appeared and whether it referred to a figure or a table. While some sentences could fall into more than one category, for the purposes of data analysis, each sentence was consistently classified into the single most representative data class. For example, a sentence that reported both previously published experimental results and novel experimental results, was classified solely as ‘Directly reports experimental results’ (e.g.: *In addition to the neuronal and intestinal expression previously detected using an acs-1 promoter-driven GFP expression construct (Kniazeva et al. 2004), we observed prominent GFP fluorescence in the somatic gonad of adults but not in that of larvae (Fig. 2A,B; Supplemental Fig. S2A–L,S,T).*) All sentences were reviewed collectively by curators to ensure accurate characterization.

#### Additional randomly selected negative sentences

For each data type, we sourced 500 additional negative sentences by randomly selecting a set of sentences from references that were manually validated negative in our curation database. Curators manually verified that these sentences were indeed negative.

#### Overall dataset

After merging all the sentences obtained as described above, we obtained an overall dataset from which we removed duplicate sentences resulting from possible overlap between the different sets.

### Sentence-level classifiers

We evaluated the performance of traditional supervised learning binary classifiers as well as Large Language Models (LLMs) in classifying sentences. For the traditional classifiers, we tested several models, including LogisticRegression, RandomForests, GradientBoosting, XGB, Multi-Layer perceptron (MLP), Support Vector Machines (SVM), K-neighbors classifiers, Stochastic Gradient Descent (SGD), and Perceptron classifiers (39–42). We converted plain text sentences into numeric vectors (sentence embedding, aka sent2vec) to be able to process them with machine learning methods. We used BioSentVec, a pretrained sent2vec model specifically designed for the biomedical literature (33), which embeds text sentences into 700-dimensional numerical vectors. Then, we fitted each model using stratified K-fold cross validation with k=5 and using hyperparameter optimization with randomized search on 100 different random configurations, for a total of 500 fits for each model. Both the hyperparameter optimization and best model selection were based on the F1-score. We calculated the average precision, recall and F1-score for the best models. We used implementations available from the Python library scikit-learn (scikit-learn.org), one of the most widely used machine learning libraries in the literature, and from the xgboost library (xgboost.readthedocs.io). For each model type we provided a range of possible hyperparameters to be selected by randomized search optimization. The code is available at https://github.com/WormBase/curation-sentence-classification.

For the LLM approach, we used BioBERT (33), available from Huggingface (huggingface.co; dmis-lab/biobert-v1.1), and GPT-4o (43). For BioBERT, we used the same model for text embedding and for classification. We fine-tuned the classification model on our specific dataset and calculated its performance using the same stratified K-fold cross validation technique used with the traditional ML models. The number of epochs for each fine-tuning phase during the K-fold cross validation was set to 5. The code is available at https://github.com/WormBase/huggingface-document-classifier.

For GPT, there are many GPT models available, such as BioGPT, but we focused on the latest models developed by OpenAI to ensure we are arguably leveraging the latest advancements in the field. Two GPT-4o models (gpt-4o-2024-08-06) were fine-tuned separately for gene expression and protein kinase activity classification. Training, validation, and testing datasets were created using a stratified multilabel split, with 70% of the data allocated for training, 15% for validation, and 15% for testing. This stratification ensured balanced representation of all curatable labels across subsets. The datasets were then converted to JSONL format as per OpenAI’s documentation (https://github.com/alliance-genome/agr_sentence_classifier). Both models were trained over 3 epochs with a batch size of 2 and a learning rate multiplier of 2. Testing of sentence datasets was conducted using custom Python code interfaced with OpenAI’s API, performing five test iterations per dataset. The evaluation workflow incorporated detailed prompts tailored to each data type for accurate classification (Supplementary File S1).

## Results

### Selection of data types

Our analysis focused on two data types curated at WormBase: gene expression and protein kinase activity. We started with these data types for several reasons. First, each of these data types is relatively straightforward for curation because they do not require curatorial interpretation, unlike for example, genetic interaction data, for which curators must sometimes interpret entire paragraphs covering multiple experiments. Secondly, sentences describing these data types can potentially be processed for automatic extraction of entities, e.g., genes, and ontology terms, such as those from the *C. elegans* Anatomy Ontology, to pre-populate curation forms. Third, we hoped to mitigate curation backlogs as exemplified by the relatively high frequency of flagging for those data types within the ACKnowledge form (e.g., as of November 2024, 336 references were flagged positive for gene expression since 2019, 20.25% of the total number of submissions). Finally, capturing the causal effects between protein kinases and their substrates is a major component of pathway modeling, and we wished to explore how sentence classification could be used to help identify experimental support for protein kinase activity annotations when creating Gene Ontology Causal Activity Models (GO-CAMs, 44).

### Gene expression curation

“Low throughput” gene expression data curation aims at extracting from the literature evidence of gene expression in a particular tissue or cell component, or during specific stages of development inferred from a small number of custom crafted experiments rather than genome-scale analyses (“High throughput”). Such gene expression experiments include reporter gene analysis, antibody staining, in situ hybridization (ISH), single-molecule fluorescent in situ hybridization (smFISH), RT-PCR, qPCR, Northern blot, Western blot analysis, and in situ hybridization chain reaction (HCR).

To make an annotation for gene expression in wild-type conditions, curators need to identify the gene product and its spatial or temporal localization. Ideally, a sentence from which a curator can make an annotation contains the gene name, a keyword indicating expression, the anatomical or cellular location of expression and/or the relevant life stage. It is worth noting that often this information is scattered across multiple adjoining sentences. Curators also record the absence of expression in specific tissues or life stages when the authors explicitly indicate the absence of the expression signal.

Notably, sentences that contain the aforementioned information may also be negative for curation, such as when authors are reporting results from previously published studies or when they are describing expression in a mutant background.

### Protein kinase activity curation

Protein kinases are enzymes that play crucial roles in regulation of biological pathways by phosphorylating specific amino acid residues to activate or inhibit the activities of their protein targets. One of the curatorial goals of WormBase and the Gene Ontology Consortium is to model such pathways using the Gene Ontology Causal Activity Model framework (GO-CAM) in which GO molecular functions (MFs) are linked to one another with causal relations, e.g. directly positively regulates [RO:0002629], from the Relations Ontology (44).

For protein kinase activity curation, we focused on sentences that report the results of biochemical experiments providing in vitro evidence for protein kinase activity, and whenever possible, the physiologically relevant substrate(s). Fully curatable experimental protein kinase activity sentences thus state the protein kinase and keywords (e.g. phosphorylation, *in vitro*) that describe the enzymatic activity and type of assay, respectively. Sentences that also state the protein kinase substrate provide additional contextual information that can be captured as part of a GO-CAM or as a standard GO annotation extension (45). We note, however, that substrate identification is not absolutely required to make a GO MF annotation. As for gene expression curation, full information for a GO MF annotation may be spread across multiple sentences.

### Characterization of data type-relevant sentences

To characterize data type-relevant sentences in depth, we analyzed the set of manually collected gene expression and protein kinase activity sentences in Supplementary Table 1, which contains additional sentence metadata compared to the randomly selected sentences in our dataset (see the Methods Section for more information on the sentence collection process). We performed this analysis because, from our curatorial experience, we have observed that there is a broad range of data type-relevant sentences in the literature and we wanted to understand the differences in those sentences in the context of curation tasks. Our goal was to define classes that could, in the future, be used for automatic sentence classification and thus aid in fact extraction. Specifically, while some data type-relevant sentences concisely report experimental results in their entirety, it is also the case that experimental results are described across multiple, sometimes non-contiguous, sentences. Also, some sentences may contain data type-relevant language without reporting curatable information.

We therefore characterized sentences across two main axes: 1) the type of information they contained, and 2) the completeness of the information presented for the purpose of creating an annotation. For example, “fully curatable” sentences contain mention of a gene or gene product, language that can be used to select an ontology term for an annotation (*C. elegans* Anatomy, Life Stage Ontologies and Gene Ontology Cellular Component for gene expression, and a Gene Ontology Molecular Function term for protein kinase activity) and, where applicable, language that allows a curator to correctly assign an evidence code for the annotation from the Evidence and Conclusion Ontology (e.g. ‘*in vitro* assay’) (46). “Partially curatable” sentences are missing one or more critical pieces of information, for example, the sentence mentions expression in an anatomy term but not the actual gene expressed or mentions the kinase substrate, but not the actual kinase, requiring a curator to seek additional curation details. “Related language” sentences may describe hypotheses or experimental design or contain similar language but not refer to a curatable experiment within that reference.

As shown in Table 1, our analyses led to the identification of three main classes of data type-relevant sentences (Column 1, Main sentence class) with several subclasses (Column 2, Specific subclass) represented within each main class.

**Table 1.** Summary of data type-relevant sentences. This table illustrates data type-relevant sentence classification as determined by expert biocurators. The three main relevant sentence classes are defined as: 1) sentences that directly report experimental results, with either complete or partially curatable information; 2) sentences that summarize experimental results, either complete or partial, including statements from reference section headings and/or figure titles; and 3) sentences that contain language relevant to a data type, but not necessarily used for curation, such as use of reporter fusions to assess phenotypic outcomes for the gene expression data type. Examples of sentences within each category are provided, highlighting the nuances in reporting and summarizing experimental data. A full list of sentence classes with relative examples is presented in Supplementary Table 1.

| **Main sentence class** | **Specific subclass** | **Protein kinase sentences examples** | **Gene expression sentences examples** |
| --- | --- | --- | --- |
| **Statements that directly report experimental results** | **Directly reports experimental results and is fully curatable** | When this activated PMK-1 was tested in the in vitro kinase assay, it was found to efficiently phosphorylate GST-SKN-1 (Fig. 3A, lane 2). | A transcriptional reporter for nlp-18 revealed strong expression in the intestine and three pairs of neurons: a gustatory sensory neuron (ASI); a mechanical sensory neuron (FLP); and an interneuron (RIM; Figures 2C and S2B). |
| | **Directly reports experimental results and is partially curatable** | When the immunoprecipitates were incubated with [γ-32P]ATP, SEK-1(K79R) became phosphorylated (Figure 3B). | We observed expression in five additional neurons close to the nerve ring, including possibly M3, AWB, and ASJ, and three neurons in the tail (Figure 1E). |
| **Summary statements** | **Summary statement of experimental results in reference and is fully curatable** | Taken together, our results demonstrate that UNC-43 is a calcium-dependent kinase capable of phosphorylating CaMKII substrates. | In conclusion, gmap1 and gmap-2/3 genes are both expressed in epithelial cells although with non-overlapping expression patterns. |
|  | **Summary statement of experimental results in reference and is partially curatable** | The protein was shown to phosphorylate a model substrate and to undergo autophosphorylation. | Expression pattern of the acox-1.1 gene. |
|  | **Summary statement of previously published experimental results** | Phosphorylation of many proteins by GSK-3 first requires that a serine or threonine four residues C-terminal be phosphorylated by a priming kinase (Biondi and Nebreda, 2003; Fiol et al., 1987). | abts-1 is expressed in HSNs from the L4 larval stage onward (Shinkai et al. 2018). |
| **Other language relevant sentences** | **Experiment was performed; experimental hypothesis or rationale** | To investigate whether MBK-2 has a direct role in OMA-1 degradation, we began by determining whether MBK-2 can phosphorylate OMA-1 in vitro. | Due to the structural and functional similarities between GPM6A, M6 and NMGP-1 and because proteolipid proteins are expressed in neurons, we hypothesized that NMGP-1 was expressed in the nervous system of the worm. |
|  | **Experimental setup** | Catalytic activity of DKF-1 was quantified by measuring incorporation of <sup>32</sup>P radioactivity from [-<sup>32</sup>P]ATP into Syntide-2 peptide substrate (Calbiochem). | Dissected gonads of adult worms were prepared for RNA in situ hybridization and processed according to the protocol described in Lee et al. (2006) (Lee and Schedl 2006). |
|  | **Related experimental results** | Critical residues for DYRK2 phosphorylation are shown in blue and red. *: the residue phosphorylated by DYRK2 in eIF2B and corresponding residues in OMA-1 and OMA-2. | As reported above, we observed that the expression and subcellular localization of RAB-11 and SYN-4 in acs-1(RNAi)C17ISO embryos were not different from wild type. |
|  | **Contains overlapping keywords with the curated data type** | An important question that remains is whether CDK-1 targets WRM-1 in a signal-dependent manner—for example, by only phosphorylating a subpopulation of WRM-1 on the membrane proximal to the P2/ EMS contact site. | Expression of CKR-1 in AIB robustly rescued ckr-1 mutant animal's escape defects (Figure 4C). |

Using the data type-relevant sentences in Supplementary Table 1 we performed a UMAP analysis on the sentence embeddings obtained with BioSentVec to see if: 1) the data type-relevant sentences were clearly separable from negative sentences that do not contain relevant language, and 2) if the sentences in the same data type-relevant sentence class cluster with each other. For this clustering analysis, we included 500 randomly selected negative sentences taken from 15 random references manually validated negative for each data type (see Methods Section for more details). As shown in Figure 1, the negative sentences (i.e. ‘does not contain language’) are separated from the data type-relevant sentences. This suggests that an automated classifier may be used to separate negative sentences from the data type-relevant classes. Resolution of individual data type-relevant sentence classes in the 2D map is not as distinct, but in some cases, e.g. the ‘Summary statements’ and the ‘Experimental set up statements’ for protein kinase activity, suggests that there are defining features in each class that could be exploited for more granular classification, given the much higher dimensionality of the original data compared to the 2D visualization obtained with UMAP. Current results demonstrate potential for further subclass separation with larger training sets.

**Figure 1.**
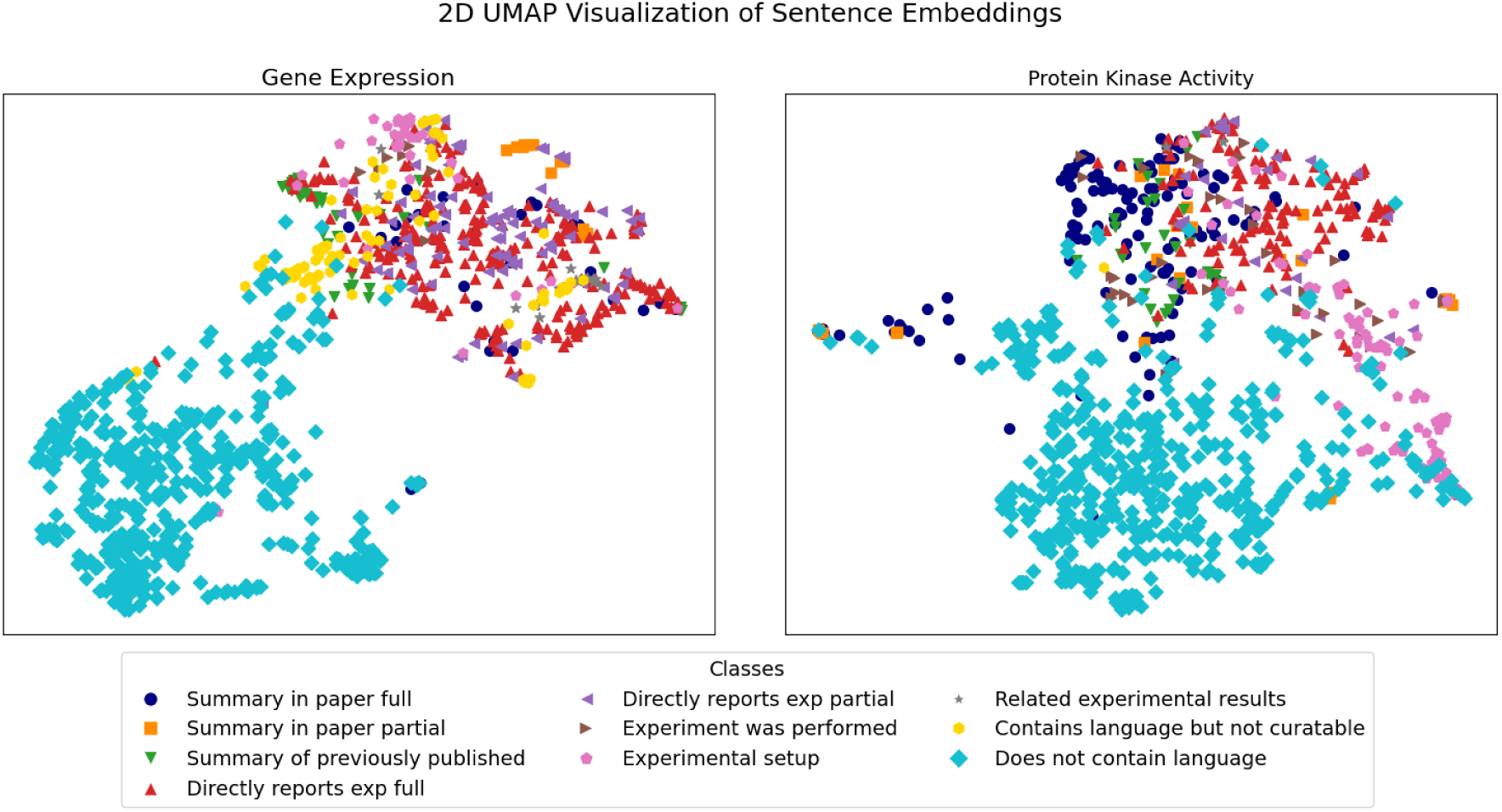
UMAP visualization of gene expression sentence embeddings (left panel) and protein kinase activity (right panel). A detailed description of the sentence classes is reported in Table 1.

**Figure 2:**
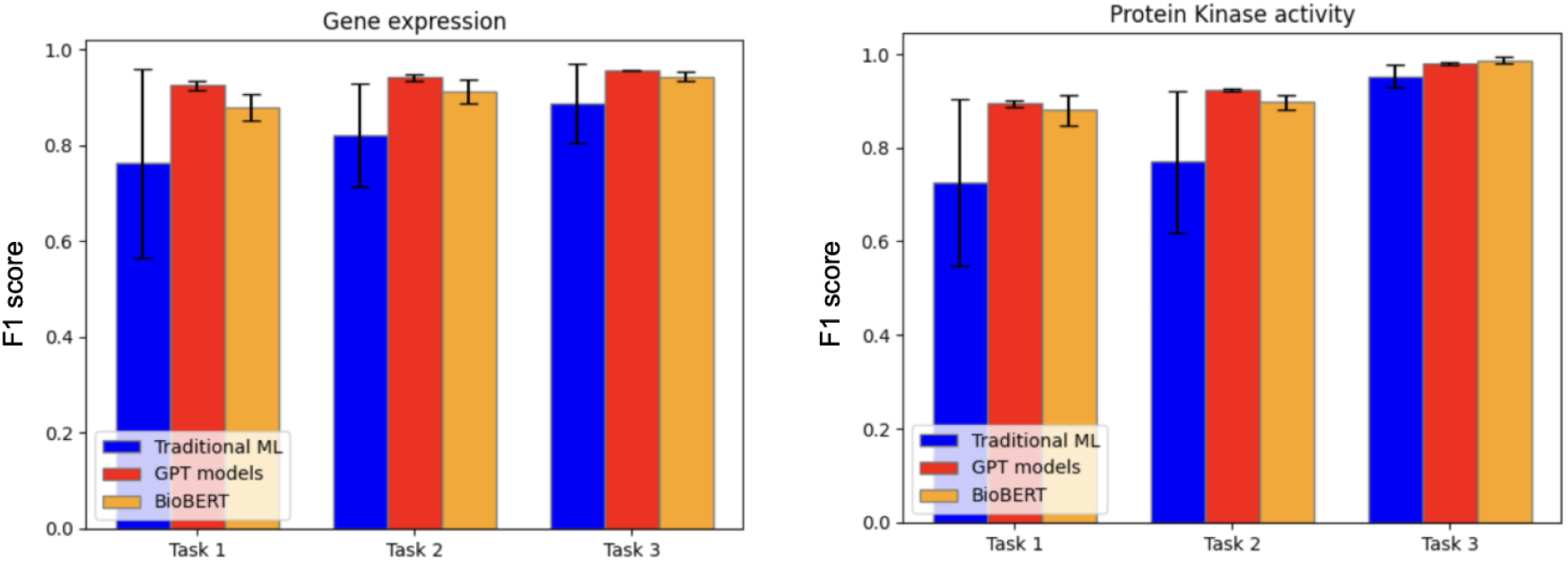
Barcharts representing the average F1 score for the different models tested for the three curation tasks.

### Curation tasks for sentence classification

Our sentence analysis illustrated that a range of sentence classes can be relevant for curation. To integrate this information into curatorial workflows, we focused on three classification tasks: i) identifying fully curatable sentences, ii) identifying partially curatable sentences, and iii) identifying sentences with related language. Table 2 presents the relevant sentence classes for each classification task along with potential use cases or applications for sentences within each class. For example, the identification of fully curatable sentences could be used to automatically create annotations or pre-populate fields in curation forms. Partially curatable sentences could be used individually, or collectively, to pre-populate curation forms or serve as curatorial aids for creating a complete annotation. Related language sentences could be used to retrieve experimental or methodological details, or for alerting curators to related experiments that warrant further review of the reference for potential curation.

**Table 2.** Task-centric sentence classification. The values in Column 1 (“Datasets composition”) indicate the task and what classes were used as positive (P) and (N) negative for that task (T). For example, T1P means Task 1 Positive). Fully curatable sentences supply all information and could be used in stand-alone curation. Partially curatable sentences would require additional text to make an annotation. Related language sentences provide hints at relevant experiments but would require additional text to confirm their presence and/or outcome.

| **Datasets composition** |  |  | **Sentences class** | **Types of sentences in this class** | **Possible use cases** |
| --- | --- | --- | --- | --- | --- |
| **T1P** | **T2P** | **T3P** | **Class A: Fully curatable** | Sentences containing all the relevant information for an annotation:<br>- Statements directly reporting experimental results and fully curatable<br>- Summary statements of experimental results in reference and fully curatable | - identify single sentences from which an annotation might be extracted automatically<br>- present “fully curatable” sentences to the authors in the ACKnowledge form and to biocurators in curation tools to aid them in creating an annotation |
| **T1N** |  |  | **Class B: Partially curatable** | Sentences containing partial relevant information for an annotation:<br>- Statements directly reporting experimental results and partially curatable<br>- Summary statements of experimental results in reference and partially curatable | - identify a set of sentences that, collectively, could be used to automatically extract an annotation<br>- present “curation relevant” sentences to the authors in the ACKnowledge form and to biocurators in curation tools to aid them in creating an annotation |
|  | **T2N** |  | **Class C: Related language** | Sentences containing related terms and phrases but not necessarily containing curatable information:<br>- Summary statement of previously published experimental results<br>- Experiment was performed; experimental hypothesis or rational<br>- Experimental setup<br>- Related experimental result<br>- Contains overlapping keywords with the curated data type | - identify additional sentences that may provide curation details (e.g. the antibody used for immunohistochemistry, the construct details for reporter gene analysis, ..)<br>- identify any sentences related to that data type that may help guide curators or readers to potentially relevant work |
|  |  | **T3N** | **Negative** | Sentences that do not contain data type-related language |  |

### Sentence classifiers - traditional ML classifiers and LLMs

To train and finetune automated sentence classifiers for the three tasks defined above, we used the complete dataset (Supplementary Table 2 - ‘Complete Sentence Datasets’), which includes data type-relevant sentences manually collected and randomly selected, as well as randomly selected negative sentences. The sentence count for each task in the dataset is summarized in Table 3. The numbers of sentences are similar for both data types. As we further discuss in the Discussion and Conclusion Section, the order of magnitude of positive sentences is sufficient to obtain significant results for all tasks for both data types, but it may be on the low side for some of the evaluated models, especially for Tasks 1 and 2.

**Table 3:**
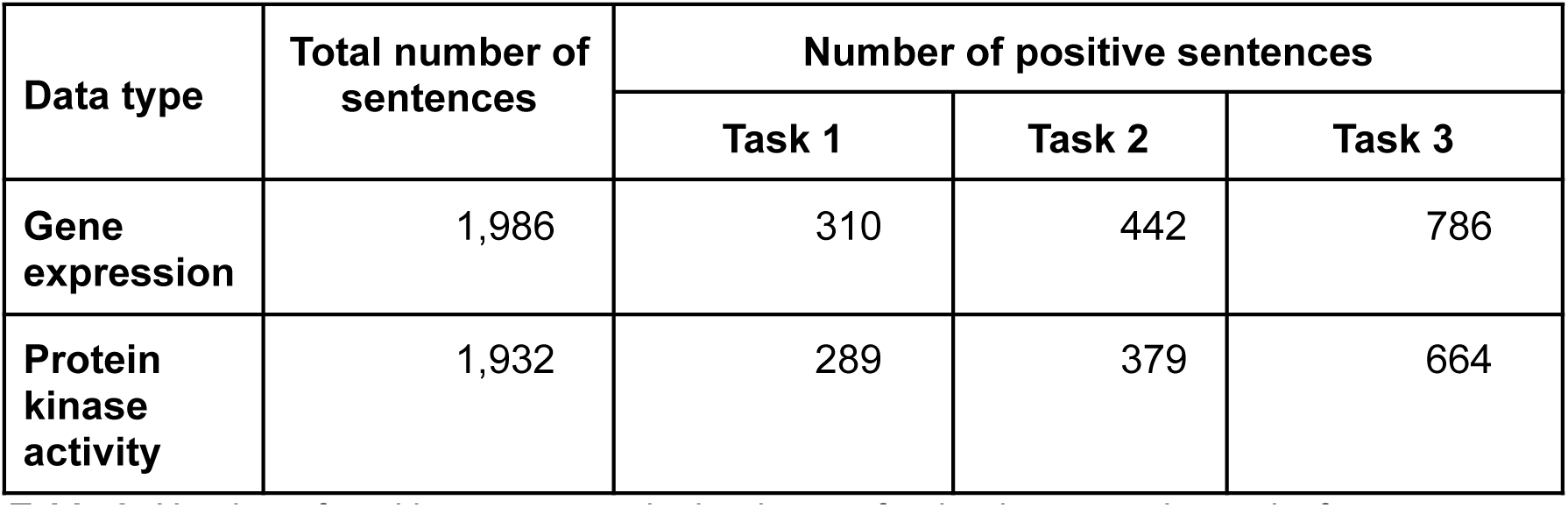
Number of positive sentences in the dataset for the three curation tasks for sentence classification.

We evaluated both traditional machine learning models and LLMs such as GPT and BERT. For the traditional models, we determined the best-performing one based on the F1 score, as described in the Methods Section. The average precision, recall, and F1 score calculated on the five folds from cross validation (as described in the methods section), for each evaluated model, data type, and task are reported in Table 4. A bar chart representation of the average F1 score is depicted in Figure 2. For traditional ML classifiers, only the best model, selected through F1 score-based model selection, is reported in Table 4.

**Table 4:** Average and standard deviation for precision, recall, and F1 score of the classifiers for gene expression and protein kinase activity for the three tasks.

| Data type | Model Type | Avg. Precision | Avg. Recall | Avg. F1 score |
| --- | --- | --- | --- | --- |
| **Task 1** |  |  |  |  |
| **Gene expression** | Traditional (SVM) | 0.718 ± 0.258 | 0.9 ± 0.031 | 0.762 ± 0.196 |
|  | GPT-4o | **0.867 ± 0.012** | **0.991 ± 0.012** | **0.925 ± 0.009** |
| | BioBERT | 0.845 ± 0.016 | 0.916 ± 0.055 | 0.879 ± 0.028 |
| **Protein kinase activity** | Traditional (SVM) | 0.688 ± 0.254 | 0.878 ± 0.090 | 0.726 ± 0.178 |
| | GPT-4o | 0.837 ± 0.008 | **0.959 ± 0.01** | **0.894 ± 0.007** |
| | BioBERT | **0.865 ± 0.078** | 0.903 ± 0.044 | 0.880 ± 0.033 |
| **Task 2** |  |  |  |  |
| **Gene expression** | Traditional (MLP) | 0.844 ± 0.195 | 0.843 ± 0.087 | 0.822 ± 0.107 |
| | GPT-4o | **0.951 ± 0.007** | 0.933 ± 0.008 | **0.942 ± 0.007** |
| | BioBERT | 0.888 ± 0.036 | **0.939 ± 0.034** | 0.912 ± 0.025 |
| **Protein kinase activity** | Traditional (Logistic) | 0.703 ± 0.221 | 0.915 ± 0.040 | 0.770 ± 0.152 |
| | GPT-4o | 0.886 ± 0.006 | **0.966 ± 0.00** | **0.924 ± 0.003** |
| | BioBERT | **0.889 ± 0.037** | 0.908 ± 0.027 | 0.897 ± 0.016 |
| **Task 3** |  |  |  |  |
| **Gene expression** | Traditional (SVM) | 0.943 ± 0.031 | 0.855 ± 0.147 | 0.888 ± 0.083 |
|  | GPT-4o | **0.966 ± 0.00** | **0.949 ± 0.00** | **0.957 ± 0.00** |
| | BioBERT | $0.948 \pm 0.016$ | $0.940 \pm 0.025$ | $0.944 \pm 0.011$ |
| **Protein kinase activity** | Traditional (SVM) | $0.966 \pm 0.022$ | $0.941 \pm 0.055$ | $0.952 \pm 0.024$ |
| | GPT-4o | $0.96 \pm 0.008$ | **$1 \pm 0.00$** | $0.98 \pm 0.004$ |
| | BioBERT | **$0.982 \pm 0.010$** | $0.991 \pm 0.009$ | **$0.987 \pm 0.008$** |

Across all tasks, GPT-4o and BioBERT showed strong performances, while traditional machine learning methods generally underperformed relative to these more advanced architectures. It is worth noting that the statistical variation of traditional ML models is rather high, at least for the first two tasks. This observation indicates that the datasets may not be large enough for these tasks. Nonetheless, the current results already give an indication of their performances. The low standard deviation of GPT and BioBERT show that these models have stable performances on different subsets of the dataset, and this is an indication that the amount of sentences used for finetuning is sufficient.

### Finding the right size for fine-tuning datasets: degradation analysis

We performed a detailed analysis to assess the minimum amount of fine-tuning data required to achieve reliable performance while minimizing the time investment of curators. This analysis is relevant, for example, to add another data type to the classification analysis, or expand the classification for that data type to other organisms. Given that collecting and curating these data demands significant curator time, our goal was to evaluate the performance drop-off at each reduction level. By doing so, we aimed to identify the minimum amount of fine-tuning data necessary for reliable performance without significantly sacrificing accuracy, thereby optimizing the trade-off between curator effort and classifier effectiveness. Given the better performances provided by BioBERT compared to traditional ML models, and the fact that it is currently less expensive to finetune than OpenAI GPT models, we decided to take BioBERT as the reference model for this analysis.

To perform the analysis, we started with the full dataset (100%) and progressively reduced the size in 10% increments, down to 10% (Figure 3). This approach allowed us to monitor how precision, recall, and F1 score were impacted as the available data decreased.

**Figure 3.**
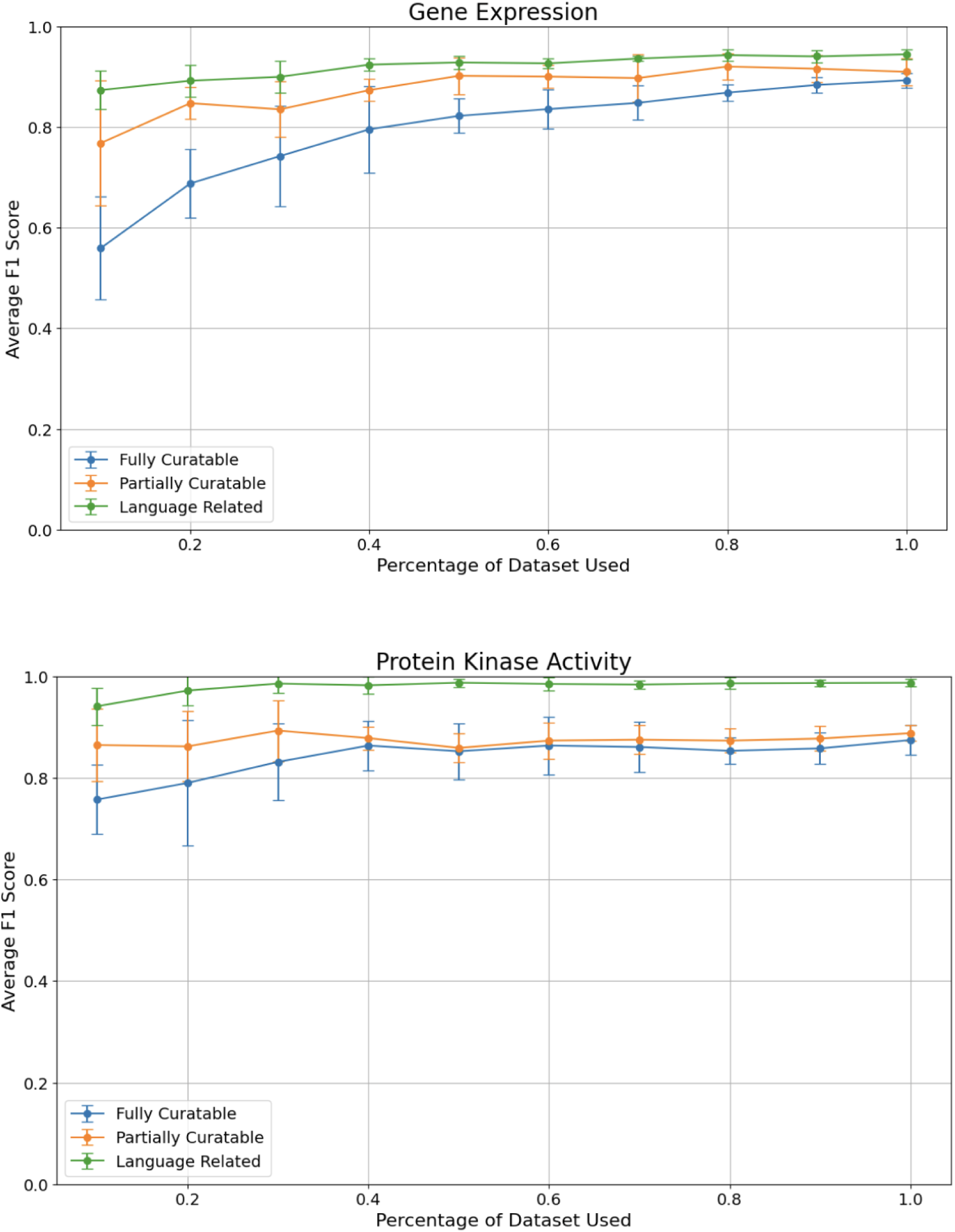
Comparison of average F1 scores across different dataset sizes for the three curation tasks: Task 1 (fully curatable), Task 2 (partially curatable), and Task 3 (language related) for the gene expression and protein kinase activity datasets.

To identify fully curatable sentences, using around 40% of our original dataset for protein kinase activity is sufficient for optimal performance. For gene expression, even though the graph does not clearly show a plateau, the gain in F1 score is marginal after 70% of the dataset. Partially curatable sentences require even less data, as low as 10% for protein kinase activity, and around 50% of the dataset for gene expression. The classification models seem to be able to correctly identify language related sentences with a small set of fine-tuning sentences for both data types, but provide the best performances after about 30%-40% of our dataset. We note that even though the standard deviation is large with a small sample size, it reduces as the number of sentences increases.

These results suggest that while a significant portion of our dataset is needed to achieve high performance, there is a point of diminishing returns where additional data does not significantly improve the model. These thresholds can help guide curators in deciding the minimum amount of fine-tuning data necessary for effective model training while balancing the time and resources required for data curation.

## Discussion and conclusions

Manual curation of the biomedical literature provides detailed, high quality annotations for biological knowledgebases and the scientific communities that rely on them daily for research support. However, this process is labor intensive and the current trend of decreased funding for knowledgebases makes it imperative that manual biocuration is augmented with high-performing machine learning methods.

In this work, we explored, for the purpose of developing automated or semi-automated curation pipelines, the manual biocuration process for two key data types, gene expression and protein kinase activity, curated by WormBase and other knowledgebases. To this end, we collected a variety of sentences used by biocurators to support detailed annotation and characterized the amount and type of information they contained. This characterization allowed us to define three sentence classification tasks for the identification of: i) fully curatable sentences; ii) partially curatable sentences; and iii) language-related sentences.

We evaluated multiple ML models to automate the classification of sentences according to these defined tasks. Among them, GPT and BioBERT models show the best average performances with very low standard deviation, even with a limited number of sentences used for fine tuning.

For implementation into biocuration workflows, we will consider the relative merits of each approach. Traditional machine learning classifiers, while showing lower performances, still achieved medium to high scores, indicating their potential utility in various curatorial tasks, especially when optimizing resource use and supporting real-time applications is critical.

BERT models tend to be slower in both training and prediction phases compared to traditional ML models, often requiring Graphics Processing Units (GPU) for efficient computation. This can become a limiting factor, especially for real-time applications or when working on very large datasets.

Large GPT models can be expensive to train and deploy, often requiring significant computational resources or high cost through proprietary APIs. On the other hand, they are very flexible models that can be used for a wide range of tasks in various applications.

Our analysis highlights the value of having well-defined tasks and high quality datasets as a baseline for training domain-specific models, such as BioBERT, or fine tuning large language models. All in all, our results indicate that it is possible to accurately classify sentences according to curatorial need, and suggest that implementing these classifiers into existing biocuration workflows has the potential to be of significant benefit to biocuration.

### Limitations

Our current datasets, consisting of approximately 2,000 sentences each, might be on the lower end of the recommended dataset size for traditional machine learning models. A common rule of thumb suggests having at least ten times as many data points as there are features in the dataset (47). Given that our sentence embeddings generated with Word2Vec result in around 700 dimensions per embedding vector, we would ideally need at least 7,000 sentences to achieve significant results. The current dataset size might therefore limit the model’s ability to generalize well and could lead to overfitting. This is reflected in the higher standard deviation in the precision, recall and F-score for traditional ML models in Table 3, especially for task 1 and 2, which have less positive sentences. On the other hand, GPT and BERT models do not necessarily need large datasets for fine tuning, due to the pre-trained nature of the models.

In the current study, we focused on analyzing two specific data types: gene expression and protein kinase activity, given their importance in WormBase curation. However, we recognize the importance of expanding our analysis to encompass a broader range of data types some of which may not be as straightforward for curation as gene expression and protein kinase activity. Therefore, we are committed to extending our research to include additional data types in future studies, and our degradation analysis can serve as a guide for minimum training sample size.

Future work to enhance the extraction of curatable information from scientific articles will focus on developing hybrid models that combine sentence-level classification with contextual information from full-text analysis, for example the location of the sentence within the article, or the content of surrounding sentences. Such models could leverage the simplicity and manageability of sentence classifiers while incorporating the contextual richness of full-text analysis. We will also leverage NLP techniques such as Named Entity Recognition (NER) to improve classification accuracy. The rapid improvements of LLMs also hold promise for high quality fact extraction and we will explore using them for suggesting annotations.

By expanding the scope of our analysis and incorporating these advanced techniques, we aim to achieve a more comprehensive and accurate extraction of curatable information from scientific literature.

As mentioned before, we will also explore ways to more rapidly create sentence datasets. One possible approach is to facilitate sentence identification and classification as part of the curatorial process. To this end, the new curation tools being developed by the Alliance of Genome Resources provide an opportunity to integrate this functionality from the outset into a curatorial framework that will be used by multiple model organism curators. Given the high recall shown by GPT models, another approach is to employ these AI models, combined with prompt engineering, to more rapidly identify and characterize curation-relevant sentences for new data types. For all approaches, we hope to systematically test the lower limits of training data to obtain the highest performance with minimum manual effort.

Lastly, to maximize the utility of the classifiers, we will integrate their output into existing curation tools such as the ACKnowledge community curation, and professional biocuration, platforms to aid in fact extraction.

In conclusion, our study demonstrates that sentence classification using advanced machine learning models, such as BioBERT and GPT, has the potential to enhance the efficiency and accuracy of biocuration. By leveraging these insights, we aim to streamline the curation process and improve the quality of annotations in biomedical knowledgebases, ultimately supporting the scientific community more effectively.

## Author Contributions

Daniela Raciti - Conceptualization, Data curation, Funding acquisition, Investigation, Validation, Writing – original draft, Writing – review & editing

Kimberly Van Auken - Conceptualization, Data curation, Funding acquisition, Investigation, Validation, Writing – original draft, Writing – review & editing Valerio Arnaboldi - Conceptualization, Formal Analysis, Funding acquisition, Investigation, Methodology, Software, Writing – original draft, Writing – review & editing

Christopher J. Tabone - Formal Analysis, Methodology, Software, Writing – original draft, Writing – review & editing

Hans-Michael Muller - Formal Analysis, Investigation, Methodology, Software, Writing – review & editing

Paul W. Sternberg - Funding acquisition, Supervision, Writing – review & editing

## Acknowledgements

We thank Chris Grove, Ranjana Kishore, and Pengyuan Li for critical feedback on this work. Their insightful comments and constructive suggestions greatly improved the quality and clarity of this publication.

