## Supplementary Table 1 Caption for "Characterization and automated classification of sentences in the biomedical literature: a case study for biocuration of gene expression and protein kinase activity"

### Spreadsheet Columns - general description

**Publication year** - the year in which the reference was published.
**Journal** - the National Library of Medicine abbreviation for the journal in which the reference was published.
**PMID** - the PubMed identifier for the reference.
**Reference section** - the part of the reference in which the sentence was found. Values for this field were recorded as they are written in the reference, e.g. using case-sensitivity.
**Location within the reference section** - the specific location in a reference where the sentence is found, e.g. in a paragraph, section heading, or figure legend.
**Text refers to figure and/or table (could be other than Fig or Table, e.g. arrowhead, panel, lane)** - Yes or No value depending on whether the sentence references a figure or table within the reference. In addition to the words 'figure' or 'table' (or variations thereof), sentences that referred to a lane, panel, arrow, or arrowhead, etc., were also recorded as positive in this field. Sentences in figure legends that do not refer to a panel, e.g. (A), or do not contain words such as lane, panel, arrow, or arrowhead, etc., cannot be distinguished from sentences in the main body of the paper and thus were marked No in this column.
**Sentence** - the text of the selected sentence.
**Types of statements:**
**Summary statement of experimental results in reference and is fully curatable** - sentences that provide a concise summary of experimental results present in the reference with all information necessary for an annotation.
**Summary statement of experimental results in reference and is partially curatable** - sentences that provide a concise summary of experimental results present in the reference but are missing at least one key aspect for an annotation.
**Summary statement of previously published experimental results** - sentences that provide a concise summary of experimental results reported in another, cited reference. These sentences often, but not always, explicitly refer to a citation, e.g. Cai et al., 2014, or a number for the cited reference, such as (25).
**Directly reports experimental results and is fully curatable** - sentences that report the actual result or observation of the experiment and have all the information necessary for an annotation. These sentences are typically found in the results section of references and often contain more experimental detail than the more concisely written summary statements.
**Directly reports experimental results and is partially curatable** - sentences that report the actual result or observation of the experiment and are missing one or more key elements of information necessary for an annotation. These sentences are typically found in the results section of papers and, while missing some information, often contain more experimental detail than the more concisely written summary statements.
**Experiment was performed; experimental hypothesis or rationale** - sentences that state that a particular type of experiment is performed. These sentences often describe the rationale for an experiment or the hypothesis being tested.

**Experimental set up** - sentences that provide specific experimental or technical details that may indicate the type of assay or reagents used and are often used to capture relevant details during the curation process.
**Related experimental results** - sentences that describe results from additional types of experiments that suggest a relevant experiment for curation of the desired data type may also be present in the reference.
**Contains overlapping keywords with the curated data type** - sentences that contain overlapping keywords with the curated data type but for which no annotation can be made.
