## Supplementary Table 2 Caption for "Characterization and automated classification of sentences in the biomedical literature: a case study for biocuration of gene expression and protein kinase activity"

**Supplementary Table 2.** This table presents the complete dataset used for the classification analysis for gene expression and protein kinase activity. Each row represents a unique sentence, with columns detailing the three classifications (fully curatable, partially curatable, and related language), and the data source.
The source column is divided into three categories:
***Gold:*** Datatype-relevant sentences, identified through manual selection.
***1000:*** Sentences collected from random sampling, derived from 15 manually validated positive papers and 15 manually validated negative papers. All sentences were manually validated.
***Negative:*** 500 additional negative sentences randomly selected from papers that were manually validated as negative for each datatype in our curation database.
Any duplicate sentences resulting from potential overlap between different sets were removed from this dataset.
